# Structural and Practical Identifiability of Dual-input Kinetic Modeling in Dynamic PET of Liver Inflammation

**DOI:** 10.1101/458927

**Authors:** Yang Zuo, Souvik Sarkar, Michael T. Corwin, Kristin Olson, Ramsey D. Badawi, Guobao Wang

## Abstract

Dynamic ^18^F-FDG PET with tracer kinetic modeling has the potential to noninvasively evaluate human liver inflammation using the FDG blood-to-tissue transport rate *K*_1_. Accurate kinetic modeling of dynamic liver PET data and *K*_1_ quantification requires the knowledge of dual-blood input function from the hepatic artery and portal vein. While the arterial input function can be derived from the aortic region on dynamic PET images, it is difficult to extract the portal vein input function accurately from PET. The optimization-derived dual-input kinetic modeling approach has been proposed to overcome this problem by jointly estimating the portal vein input function and FDG tracer kinetics from time activity curve fitting. In this paper, we further characterize the model properties by analyzing the structural identifiability of the model parameters using the Laplace transform and practical identifiability using Monte Carlo simulation based on fourteen patient datasets. The theoretical analysis has indicated that all the kinetic parameters of the dual-input kinetic model are structurally identifiable, though subject to local solutions. The Monte Carlo simulation results have shown that FDG *K*_1_ can be estimated reliably in the whole-liver region of interest with reasonable bias, standard deviation, and high correlation between estimated and original values, indicating of practical identifiability of *K*_1_. The result has also demonstrated the correlation between *K*_1_ and histological liver inflammation scores is reliable. FDG *K*_1_ quantification by the optimization-derived dual-input kinetic model is promising for assessing liver inflammation.

## 1. Introduction

Nonalcoholic steatohepatitis (NASH) is a progressive nonalcoholic fatty liver disease (NAFLD) affecting approximately 5-10 million patients in the United States [Michelotti et al., 2013, Musso et al., 2011]. The hallmark of NASH is hepatic inflammation and injury in the setting of hepatic steatosis [Wree et al., 2013]. While invasive liver biopsy is the current gold standard in clinics, dynamic 18F-fluorodeoxyglucose (FDG) positron emission tomography (PET) with kinetic modeling has been demonstrated to be promising for assessing liver inflammation non-invasively by quantifying the FDG blood-to-tissue transport rate *K_1_* [Wang et al., 2017, Sarkar et al., 2017, Sarkar et al., 2018]. Beasue the liver recieves dual blood supplies from the hepatic artery and portal vein, accurate liver PET kinetic modeling and quantification of *K*_1_ require the knowledge of dual-blood input function (DBIF) [Wang et al., 2018, Munk et al., 2001]. Although the arterial input function can be derived from the aortic region on dynamic PET images, it is difficult to extract the portal vein input function from PET. The limited spatial resolution of PET and small anatomic size of the portal vein result in serious partial volume effects and high noise in the image-derived input function.

Traditional single-input kinetic modeling neglects the difference between the hepatic artery input function and portal vein input function, resulting in inaccuracy in kinetic parameter estimation [Brix et al., 2001, Munk et al., 2001]. Existing population-based DBIF approaches [Brix et al., 2001, Munk et al., 2001, Kudomi et al., 2009] use the model parameters pre-determined by population means that were commonly derived using arterial blood sampling in animal studies, which however can become ineffective in human studies. In contrast, the optimization-derived DBIF model [Wang et al., 2018] employs mathematical optimization to jointly estimate the parameters of DBIF and liver FDG kinetics. It directly utilizes image-derived arterial input function, requires no invasive arterial blood sampling, and is more adaptive to individual patients. With the improved kinetic modeling, the FDG blood-to-liver transport rate *K*_1_ was statistically associated with histopathologic grades of liver inflammation, while *K*_1_ by the traditional SBIF model and population-based DBIF model did not show a statistical significance [Wang et al., 2018].

Identifiability analysis is crucial for examining the stability of a kinetic model [Gunn, 1996, Mankoff et al., 2006]. It characterizes whether or not the unknown parameters of a specified model can be uniquely determined and how reliably these parameters can be estimated from noisy measurements. This concept was first brought to the field of biological system research by Bellman and Astrom [Bellman and Astrom, 1970] as an extension of studies on control systems [Kalman, 1963]. Identifiability analysis has been widely used in mathematical modeling in economy [Rothenberg, 1971], chemistry [Komorowski et al., 2011] and system biology [Raue et al., 2014] and dynamic PET [Gunn, 1996, Mankoff et al., 2006]. There are two main types of identifiabilities - structural identifiability [Bellman and Astrom, 1970, Anderson, 1983] and practical identifiability [Miao et al., 2008, Miao et al., 2011]. Structural identifiability refers to under what condition the kinetic parameters can be determined from noise-free data. Practical identifiability refers to how reliably a structurally identifiable parameter can be estimated from noisy data. Note that even if a parameter is structurally identifiable, it may not be estimated with adequate accuracy from real measurements. The analysis methods for structural identifiability include the Laplace transform [Bellman and Astrom, 1970], power series expansion [Pohjanpalo, 1978], similarity transform [Walter and Lecourtier, 1981], differential algebra [Ljung and Glad, 1994, Audoly et al., 2001, Xia and Moog, 2003] and so on [Miao et al., 2011]. The methods for practical identifiability analysis include Monte Carlo simulation [Miao et al., 2008], correlation matrix [Rodriguez-Fernandez et al., 2006a, Rodriguez-Fernandez et al., 2006b] and so on [Miao et al., 2011]. Despite its high computational cost, Monte Carlo simulation is considered as the most effective method for analyzing the practical identifiability of a model.

In dynamic PET, most of the popular kinetic models follow the first-order ordinary differential equations with linear parameters and are commonly structurally identifiable. Hence, previous identifiability studies in dynamic PET focused on practical identifiability analysis [El Fakhri et al., 2009, Mankoff et al., 1998, Muzi et al., 2006, Doot et al., 2010, Muzi et al., 2005]. For dual-input kinetic modeling, the optimization-derived DBIF model contains two additional parameters when compared with traditional SBIF and population-based DBIF model. While the new model has improved the practical correlation of FDG *K*_1_ with histology, it is still unclear if all the free parameters are identifiable. In addition, the increased number of free parameters may potentially increase variance in kinetic parameter estimation, but little is known so far on the quantitative aspects of the modeling.

In this paper, we conduct a theoretical analysis using the Laplace transform to assess the structural identifiability and conduct a Monte Carlo simulation to evaluate the practical identifiability using patient data of liver inflammation. The results from this study can be used to indicate the quantification accuracy and precision of model parameters and provide guidance for further improving kinetic modeling of dynamic liver PET data.

## 2. Structural identifiability analysis using the Laplace transform

### 2.1. Compartmental modeling by differential equations

Most compartmental models in dynamic PET imaging can be described by the following firstorder ordinary differential equations:

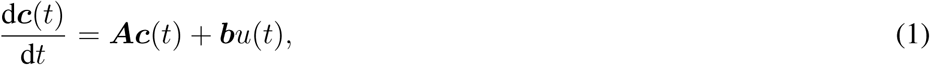

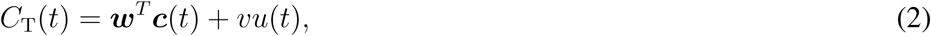

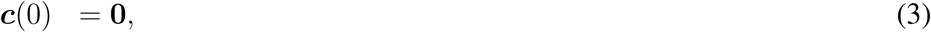

where *t* is time; ***c***(*t*) = [*c*_1_ (*t*),*c*_2_(*t*),…, *c_n_*(*t*)]*^T^* is the system states which are assumed to be zero at initial time, where *c_i_* represents the time activity of the *i*-th compartment and *n* is the number of tissue compartments; *u*(*t*) is the system input, often representing the blood input function in dynamic PET; *C_T_* (*t*) is the system output, i.e., the measured time activity curve (TAC) by PET; ***A*** is a *n* by *n* matrix, ***b*** is a *n* by 1 vector, *w* is a *n* by 1 vector and *υ* is a scalar vector. ***A, b, w*** and *υ* are composed of the kinetic parameters ***θ*** = (*θ*_1_,*θ*_2_,…, *θ_m_*) to be determined.

For a commonly used 3-compartment model (Fig. 1) such as for dynamic FDG-PET imaging, we have

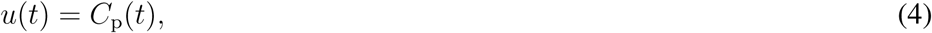

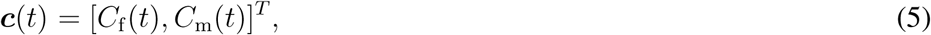

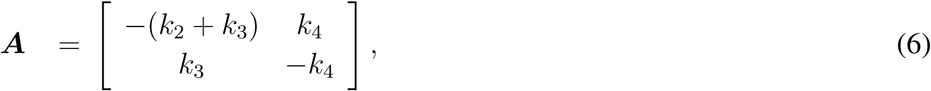

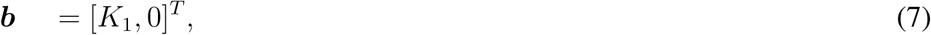

**Figure 1.**
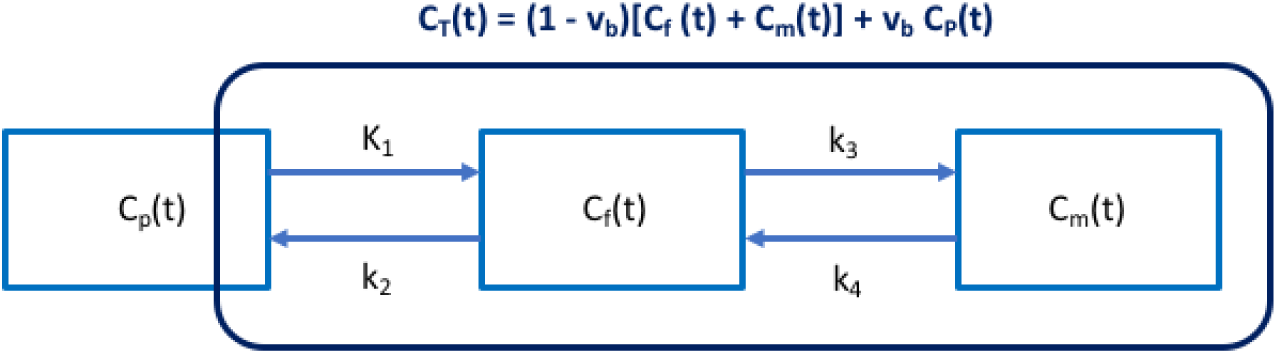
Three-compartment (*C*_p_(*t*), *C*_f_(*t*), *C*_m_(*t*)) model with single-blood input function (SBIF). *C*_T_(*t*) denotes the total activity that can be measured by PET.

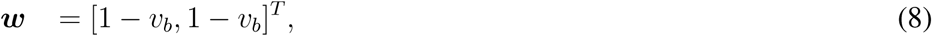

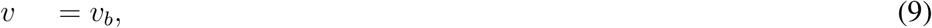

where *C_p_*(*t*) is the plasma input function; *C*_f_ (*t*) and *C*_m_(*t*) are the concentration in the free FDG and metabolized FDG compartments, respectively. The superscript “T” denotes matrix or vector transpose. ***θ*** = [*υ_b_*, *K*_1_, *k*_2_, *k*_3_, *k*_4_] with *K*_1_, *k*_2_, *k*_3_, *k*_4_ denoting the rate constants of FDG transport among compartments. *υ_b_* denotes the fractional blood volume.

The optimization-derived DBIF model [Wang et al., 2018] for analyzing dynamic liver PET data is shown in Fig. 2 and can be described using the following expressions:

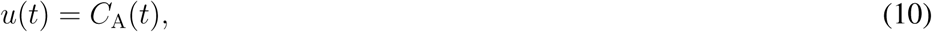

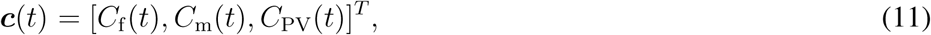

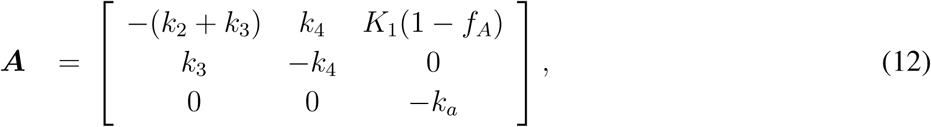

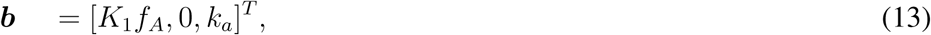

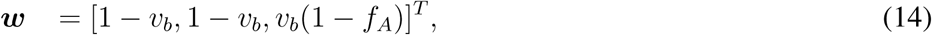

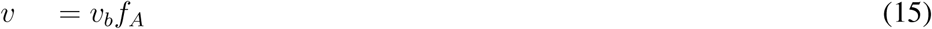

where *C*_A_(*t*) denotes the blood input function extracted from the hepatic artery; *C*_PV_(*t*) is the portal vein input function; *k_a_* is the rate constant with which FDG flows through the gastrointestinal system. *f_A_* is the fraction of hepatic artery contribution to the overall liver blood flow. The parameters to be determined are ***θ*** = [*υ_b_*, *k*_1_, *k*_2_, *k*_3_, *k*_4_, *k_a_*, *f_A_*]^*T*^.

**Figure 2.**
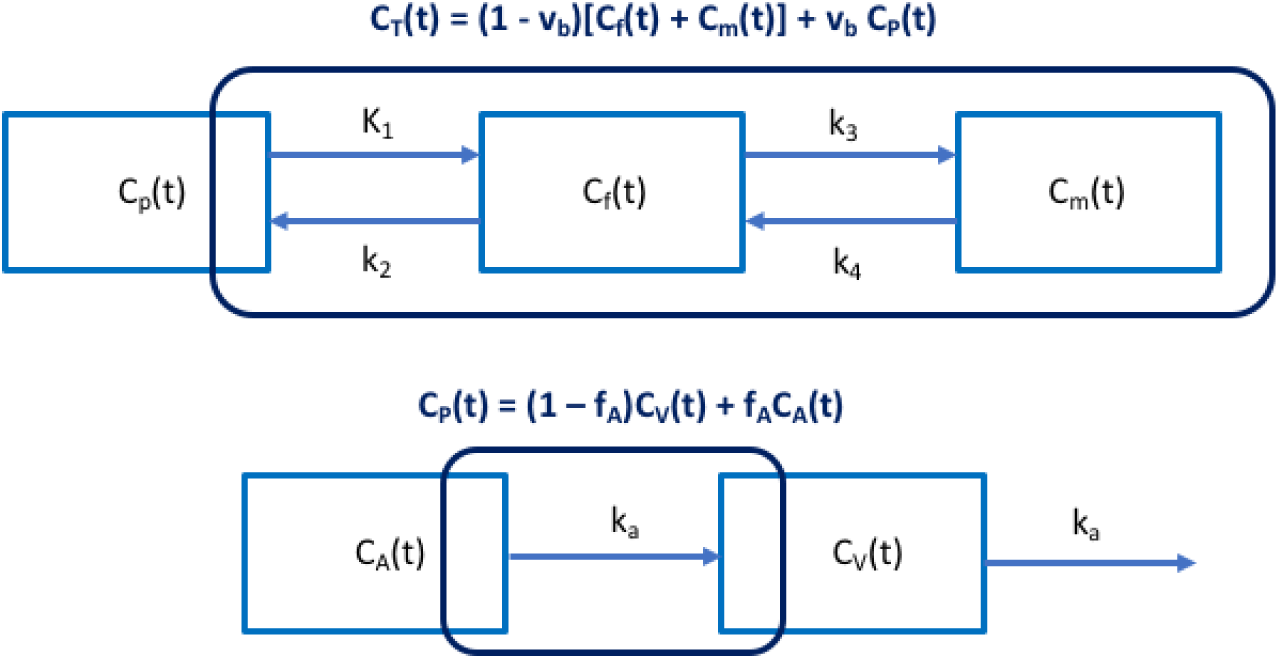
Optimization-derived dual-blood input function (DBIF) model. The FDG kinetic parameters (*υ_b_*, *K*_1_, *K*_2_, *K*_3_, *K*_4_) and dual-input parameters (*f_A_*, *k_a_*) are jointly estimated by time activity curve fitting.

### 2.2. Laplace transform for structural identifiability analysis

The Laplace transform method is a popular method in the field of system theory for analyzing differential equations [Oppenheim et al., 1996, Tsien, 1954]. After the transform, the time derivative *∂*/*∂t* becomes a multiplication of frequency *s*, thus simplifying the mathematical analysis. Taking the Laplace transform of equations (1) - (2) and making use of equation (3), one has

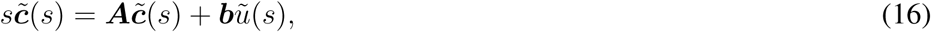

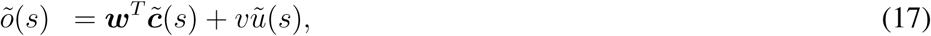

where

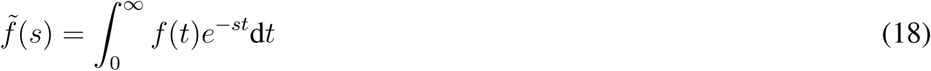

represents the Laplace transform of any function *f* in the time domain.

The system input-output relation can then be expressed as

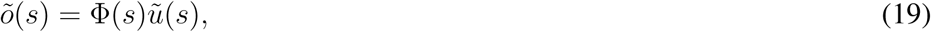

where Φ(*s*) is called the transfer function in the frequency domain,

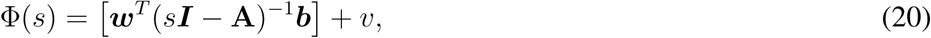

with *I* denoting the identity matrix. Φ(*s*) can be further expressed as a fractional function

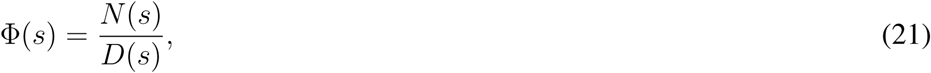

where both the numerator *N*(*s*) and denominator *D*(*s*) are a polynomial of the frequency *s*:

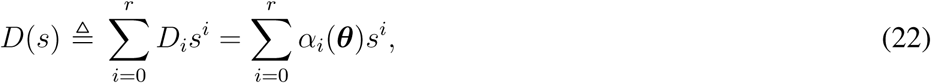

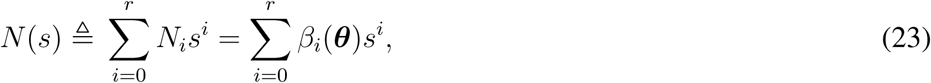

with *r* being the highest order of the polynomials of *s*. *N_i_* is the coefficient of order *i* in *N*(*s*) and *D_i_* is the coefficient of order *i* in *D*(*s*). *α_i_*(***θ***) and *β_i_*(***θ***) describe the theoretical model of *N_i_* and *D_i_* with respect to ***θ***, respectively.

The structural identifiability analysis examines if the unknown parameter set ***θ*** can be uniquely determined from 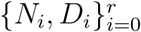 following the equation set:

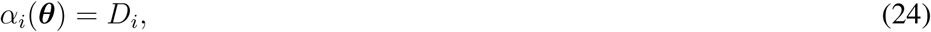

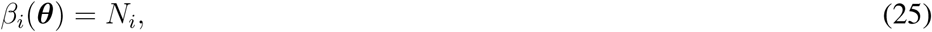

for *i* from 0 to *r*. If there are arbitrary solutions for the equations, the model structure is non-identifiable. If the equations have a unique solution for any admissible input and in the whole parameter space, the model structure is called globally identifiable. If the solution only holds unique for a neighborhood of some points ***θ***_*_ in the parameter space, the structure is then locally identifiable.

### 2.3. Structural identifiability of single-input kinetic model

Substituting equations (4) - (9) into equation (20), we have

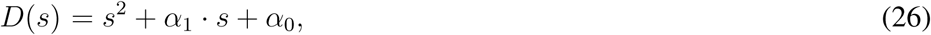

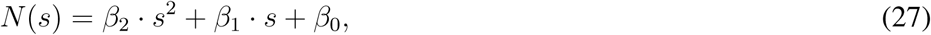

where the coefficients are defined by

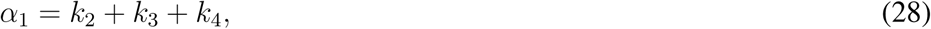

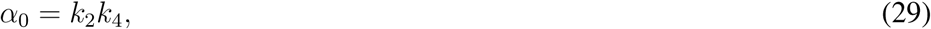

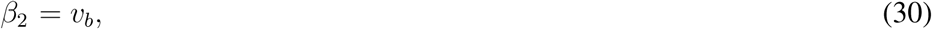

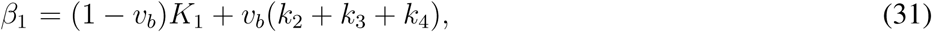

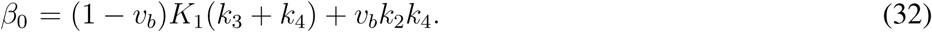

Using the equation set *α_i_* = *D_i_* and *β_i_* = *N_i_*, we can obtain a unique solution for ***θ*** after some algebraic operations:

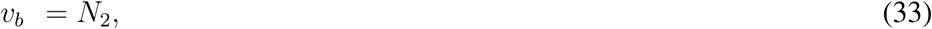

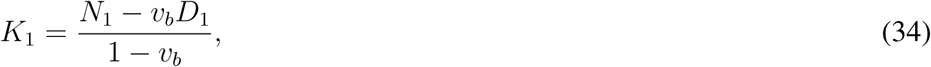

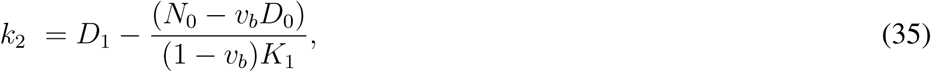

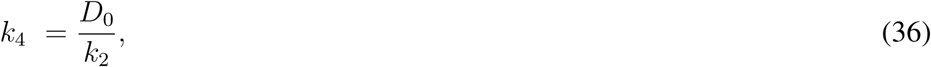

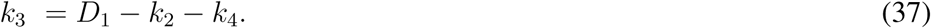

Therefore, the traditional SBIF three-compartmental model structure is globally identifiable in the parameter space.

### 2.4. Structural identifiability of dual-input kinetic modeling

For the optimization-derived DBIF model, the numerator and denominator of the transfer function Φ(*s*) are given by

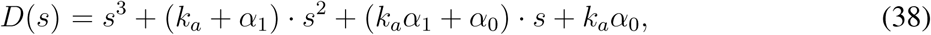

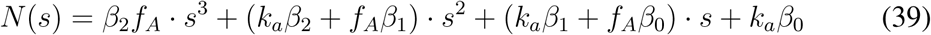

where {*α_i_*, *β_i_*} are defined by equations (28)-(32). The equation set to determine ***θ*** is:

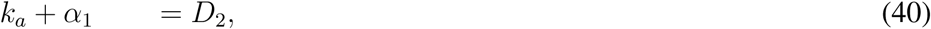

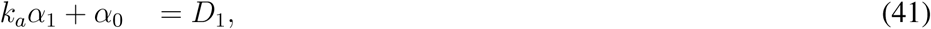

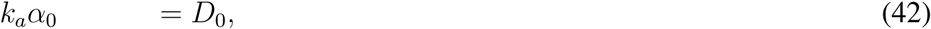

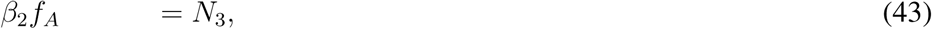

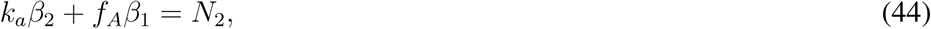

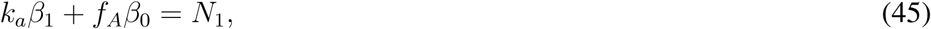

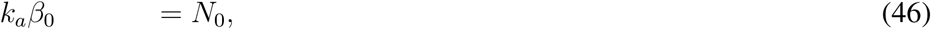

Using equations (40) - (42), we can derive

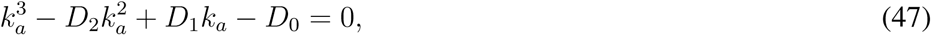

which is a cubic equation of *k_a_*. In the real parameter space, the number of roots for *k_a_* is at least 1 and at most 3. Similarly, we can derive *f_A_* using equations (43) - (46),

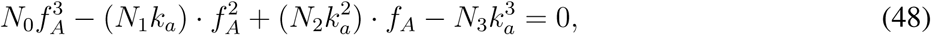

which is also a cubic equation when *k_a_* is fixed. The number of roots for *f_A_* in the non-negative parameter space is at least 1 and at most 3.

These results indicate that *k_a_* and *f_A_* are not globally identifiable because they may have multiple solutions. However, given there is at least one nonnegative root for *k_a_* and *f_A_*, they are locally identifiable. In practice, this requires a proper definition of initial estimates, lower and upper bounds for the parameters. Once *k_a_* and *f_A_* are determined, all *α_i_* and *β_i_* are determined. *K*_1_, *k*_2_, *k*_3_, *k*_4_ can then be determined from *α_i_* and *β_i_* as the same as in the single-input kinetic modeling.

## 3. Practical identifiability analysis using Monte Carlo simulation

### 3.1. Monte Carlo simulation

#### 3.1.1. Overall description

The process of the Monte Carlo simulation is described in figure 3. For each simulation, the nominal kinetic parameters ***θ***_0_ and the input function *C_A_*(*t*) were extracted from one of the human patient datasets and used to generate the noise-free liver tissue time activity curves (TAC). Independently and identically distributed noise was then added to the noise-free TAC following a defined noise model to generated *N* = 1000 realizations of noisy tissue TAC. We then fit the noisy liver tissue TACs and estimate the kinetic parameters using the optimization-derived DBIF kinetic model and the noisy aortic input function. Bias and standard deviation (SD) were calculated to assess the statistical properties of each kinetic parameter estimation. This Monte Carlo simulation was repeated for multiple patient data sets.

**Figure 3.**
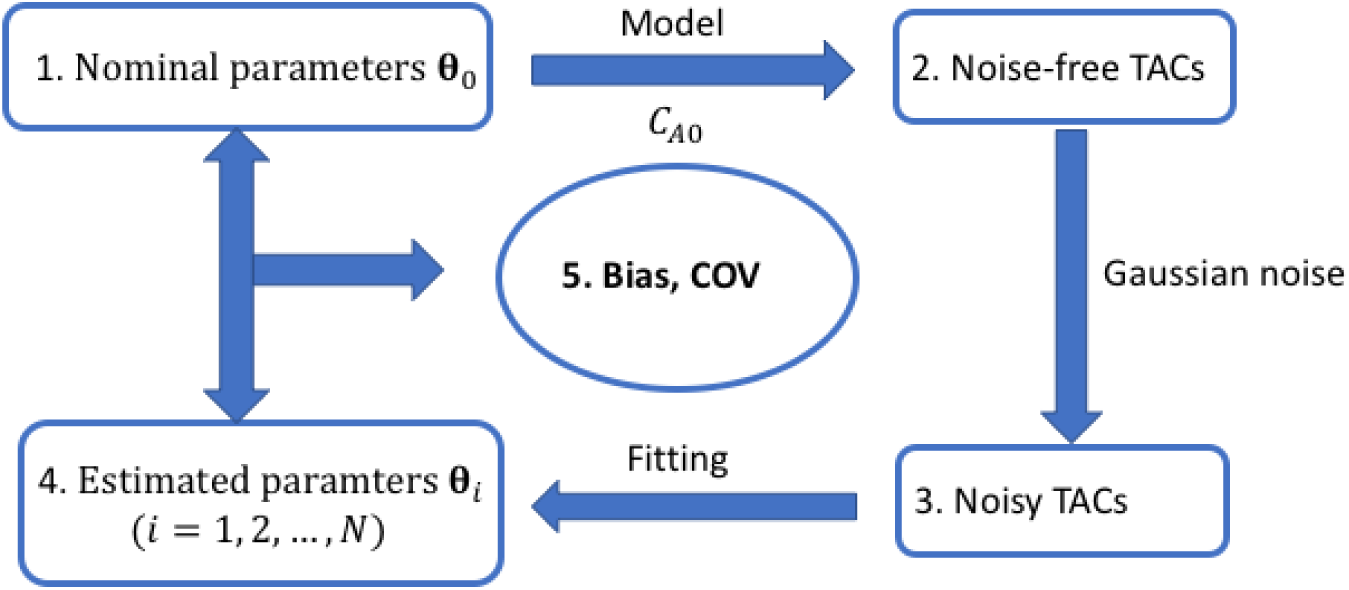
Flowchart of the practical identifiability analysis using Monte Carlo simulation.

#### 3.1.2. Human liver FDG kinetics and histological data

Fourteen patients with NAFLD were included in this study to provide nominal kinetic parameters. These patients had a liver biopsy as a part of routine clinical care or for enrollment in clinical trials. Liver biopsies were scored according to the nonalcoholic steatohepatitis clinical research network (NASH-CRN) criteria. The scores of lobular inflammation and ballooning degeneration are combined to create an overall “liver inflammation” score (range 0-5). Dynamic PET studies were performed using the GE Discovery 690 PET/CT scanner at the UC Davis Medical Center. Each patient was injected with 10 mCi ^18^F-FDG and scanned for one hour, followed by a transmission CT scan for attenuation correction at the end of PET scan. Dynamic PET data were reconstructed into 49 time frames (30 x 10s, 10 x 60s, and 9 x 300s) using the vendor software with the standard ordered subsets expectation maximization algorithm with 2 iterations and 32 subsets.

Eight spherical regions of interest (ROI), each with 25 mm in diameter, were placed on eight segments of the liver excluding the caudate lobe and avoiding any major blood vessels. An illustration is shown in Figure 4. A TAC was extracted from each liver-segment ROI. The average of these TACs was used to represent the tissue TAC in the whole-liver region. An additional volumetric ROI was placed in the descending aorta region to extract image-derived aortic input function. The optimization-derived DBIF model was used to derive the regional liver FDG kinetics at both the whole-liver ROI level and liver-segment ROI level. Hence there are a total 14 whole-liver FDG kinetic parameter sets and 112 liver-segment kinetic parameter sets from the 14 patient scans.

**Figure 4.**
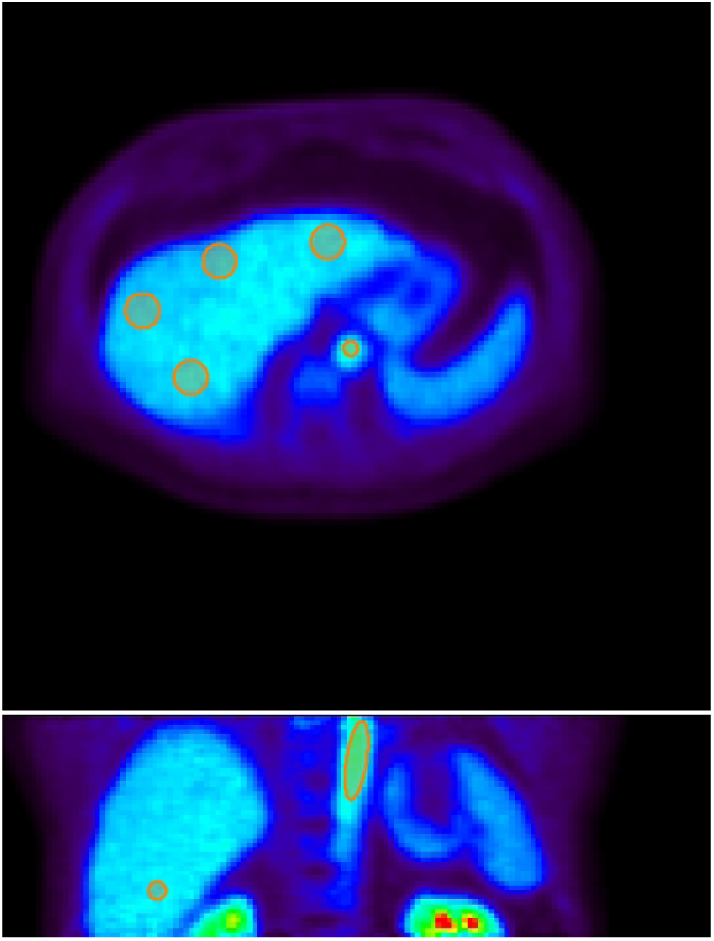
Illustration of volumetric ROIs in the liver segments and aorta in 2D planes. Top: a transverse plane showing the aortic ROI and four of eight spherical liver ROIs; Bottom: a coronal plane showing the aortic ROI and one of eight spherical liver ROIs. ROIs are overlayed on the PET image of one-hour duration. All the spherical liver ROIs are of 25 mm in diameter.

#### 3.1.3. Noise model of TACs

The reconstructed time activity in the frame *m*, *c_m_*, can be approximately modeled by an i.i.d. Gaussian distribution [Wu and Carson, 2002, Carson et al., 1993],

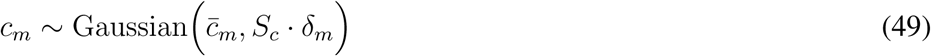

where {*c̅_m_*} denotes the noise-free TAC and *S_c_* is a scaling factor adjusting the amplitude of the unscaled standard deviation *δ_m_*,

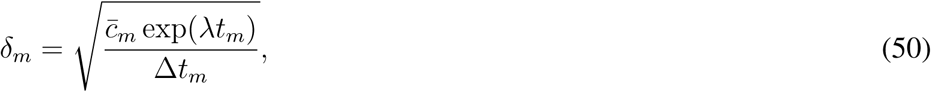

where *t_m_* is the mid-time of frame *m*, Δ*t_m_* is the scan duration of the time frame *m*, and λ = ln2/*T*_1/2_ is the decay constant of radiotracer with *T*_1/2_ (min) being the half-life. For ^18^F-FDG, *T*_1/2_ = 109.8 minutes.

Equivalently, the normalized residual difference follows a zero-mean Gaussian with the standard deviation *S_c_*:

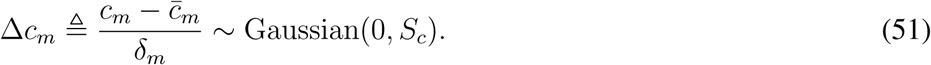

From our patient study, we have a total 14 patients × 49 frames/patient = 686 samples for Δ*c_m_* extracted at the whole-liver ROI level. The scale *S_c_* can then be determined by approximating the histogram of Δ*c_m_* using the Gaussian with the standard deviation *S_c_*. Similarly, we have a total 686 × 8 = 5488 samples to estimate *S_c_* for the noise level at the liver-segment ROI level.

### 3.2. Analysis methods

#### 3.2.1. Sensitivity analysis

The normalized sensitivity of a model TAC *C*_T_(*t*) with regard to a kinetic parameter *θ_k_* is defined by

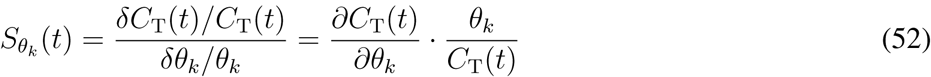

where *k* denotes the *k*th element of the kinetic parameter set ***θ*** and *∂C*_T_(*t*)/*∂θ_k_* denotes the partial derivative of *C*_T_(*t*) with respect to *θ_k_*. The normalized sensitivity function illustrates how much the model TAC would change in response to a small change *δθ_k_* in the individual parameter *θ_k_*. We evaluated the sensitivity functions for the mean of the kinetic parameters of the 14 patient datasets.

#### 3.2.2. Quantification of bias and SD of kinetic parameters

For each true kinetic parameter 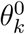, the percent bias and SD of the kinetic parameter estimate 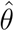 are calculated as

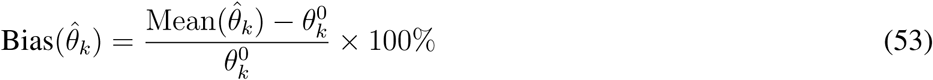

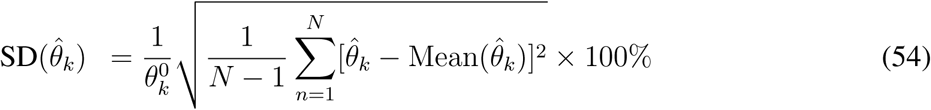

where Mean(·) represents the mean of the kinetic parameter estimates 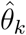, respectively.

#### 3.2.3. Comparison of different fitting options

The initial values of the kinetic parameter set [*υ_b_*, *k*_1_, *k*_2_, *k*_3_, *k*_4_, *k_a_*, *f_A_*] were set to [0.01, 1.0, 1.0, 0.01, 0.01, 1, 0.01] with lower bound [0, 0, 0, 0, 0, 0, 1, 0] and upper bound [1, 10, 10, 1, 0.1, 10, 1]. The weighting factor for the fitting was also initially set to be uniform as used in our previous study [Wang et al., 2018]. Nevertheless, our initial analysis indicates that this initialization may result in significant bias in *K*_1_ for some patient datasets. To solve this problem, we proposed two modifications to improve the fitting and *K*_1_ quantification. Instead of using a single initial value 1.0, we repeated the TAC fitting using different *K*_1_ initial values (0.5, 1.0, 1.5, 2.0, 2.5, 3.0). The one with minimum least-squares of TAC fitting was used as the optimal. This modification can reduce the effect of getting stuck at a local solution of *K*_1_. In addition, we also tested a nonuniform weighting scheme *w_m_* = Δ*t_m_* · exp (−λ · *t_m_*) versus the uniform weighting scheme *w_m_* = 1. As *K*_1_ is the major parameter of interest, these different approaches were compared for reducing the bias of *K*_1_.

#### 3.2.4. Parameter estimation accuracy over clinical range

In addition to evaluating the bias and SD for each individual kinetic data set, we also evaluated the overall performance of the model over a wide parameter range following the approach used in [Mankoff et al., 1998]. We conducted a Pearson’s linear correlation analysis to assess the closeness between 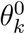 and 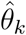 of all patients. The closer the correlation coefficient *r* is to 1, the more reliable the parameter can be estimated by the model over a wide range. In this study, we used the liver-segment kinetic parameter sets to allow a wide range of valuates to form the correlation plot.

#### 3.2.5. Variation of the correlation between FDG K_1_ and liver inflammation

Our previous study of a patient cohort of 14 patients had demonstrated that the FDG K_1_ parameter correlated with histological liver inflammation score with a statistical significance. Here we evaluate the reliability and uncertainty associated with the correlation between PET *K*_1_ and histology. This is done by repeating the estimation of 14 patient kinetic parameter sets at the whole-liver ROI level for *N* = 1000 noisy realizations using the Monte Carlo simulation study (Fig. 3). The linear correlation *r* between liver inflammation score and *K*_1_ is calculated for each realization. The bias, standard deviation and 95% confidence interval of r were then calculated to assess the reliability.

### 3.3. Results

#### 3.3.1. Sensitivity analysis

Figure 5 shows the plots of normalized sensitivity functions for different kinetic parameters in the optimization-derived DBIF model. The parameter set was the population means ***θ***̅ = [0.0185,1.0013,1.1400, 0.0149, 0.0586,1.9849, 0.0405]^*T*^. The plots of *υ_b_*, *k_a_*, *f_A_* are zoomed in for the first ten minutes for better demonstration of the differences. The sensitivity curves of *K*_1_, *k*_2_ and other vascular-related kinetic parameters *υ_b_*, *k_a_*, *f_A_* become very stable after *t* = 15 minutes, indicating the early-time data dominate the estimation of these parameters. In comparison, the curves of *k*_3_ and *k*_4_ keep increasing or decreasing in the first 60 minutes, suggesting the estimation of these two parameters needs a sufficient long scan. *K*_1_ and *k*_2_ had opposite effect on the overall uptake with greater contributions than *k*_3_ and *k*_4_. The absolute sensitivity of *K*_1_ is different from that of *k*_2_ mainly in the early time.

**Figure 5.**
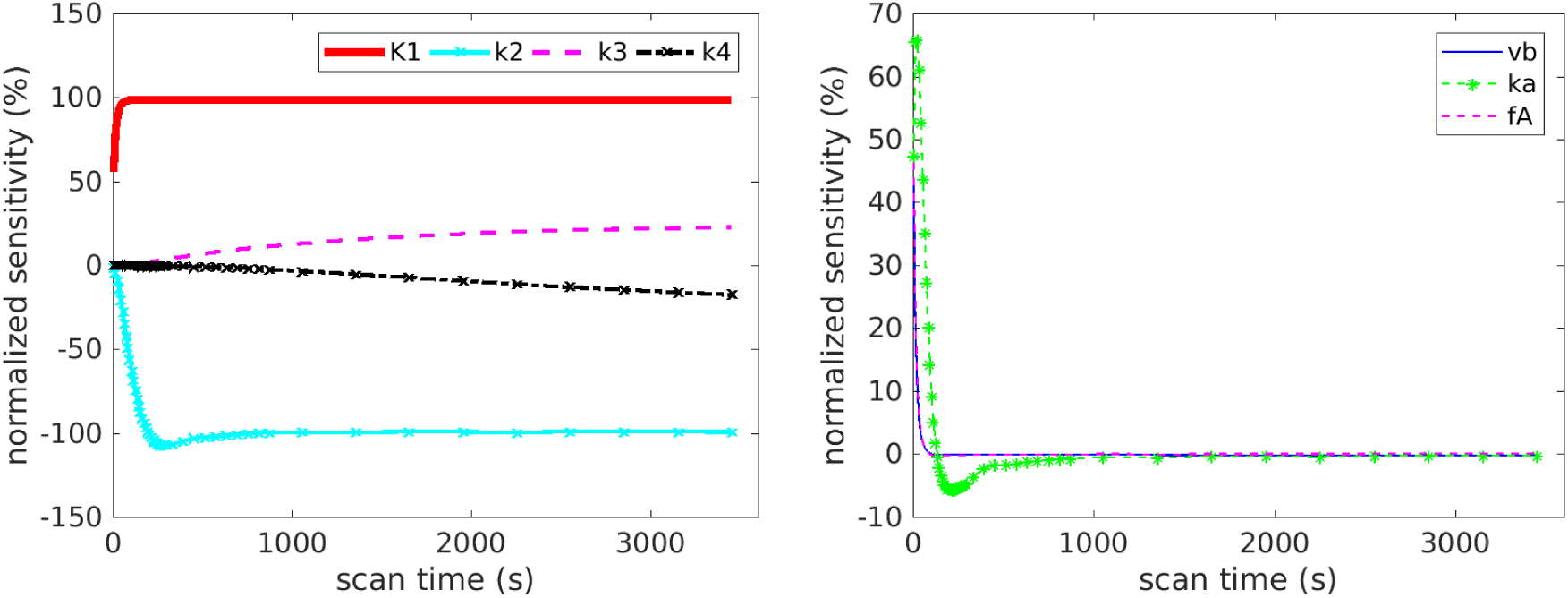
Normalized sensitivity functions representing percent change in model TAC *C*_T_(*t*) in response to the change in kinetic parameters (a) *K*_1_, *k*_2_, *k*_3_, *k*_4_ and (b) *υ_b_*,*k_a_*,*f_A_*.

The sensitivity curves of *υ_b_*,*k_a_*,*f_A_* become nearly zero after *t* =15 minutes, suggesting the late time data contribute trivial to the estimation of these parameters. The curve of *υ_b_* is almost fully overlapped with that of *f_A_*. This indicates it is difficult to differentiate them from each other. Because the curve shape of *υ_b_* or *f_A_* is different from the shape of *K*_1_, the coupled effect of *υ_b_* and *f_A_* should have a minimal effect on the estimation of *K*_1_.

#### 3.3.2. Determination of the noise model parameter

Figure 6 shows the histograms of the normalized residual error Δ*c_m_* at the whole-liver ROI level and the liver-segment ROI level. The obtained *S_c_* values are 0.3 and 0.6, respectively. The distribution of Δ*c_m_* approximately follows a Gaussian well in both cases. We therefore used these two *S_c_* values to define the noise standard deviation for the whole-liver ROI level and liver-segment level in the simulation studies. Note that the size of whole-liver ROI is 8 times that of the liver-segment ROI, which is supposed to reduce the noise standard deviation by a factor of 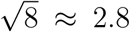. However, pixels in the liver ROI are not fully independent of each other and therefore the reduction in *S_c_* can be smaller than that by the ROI size increase.

**Figure 6.**
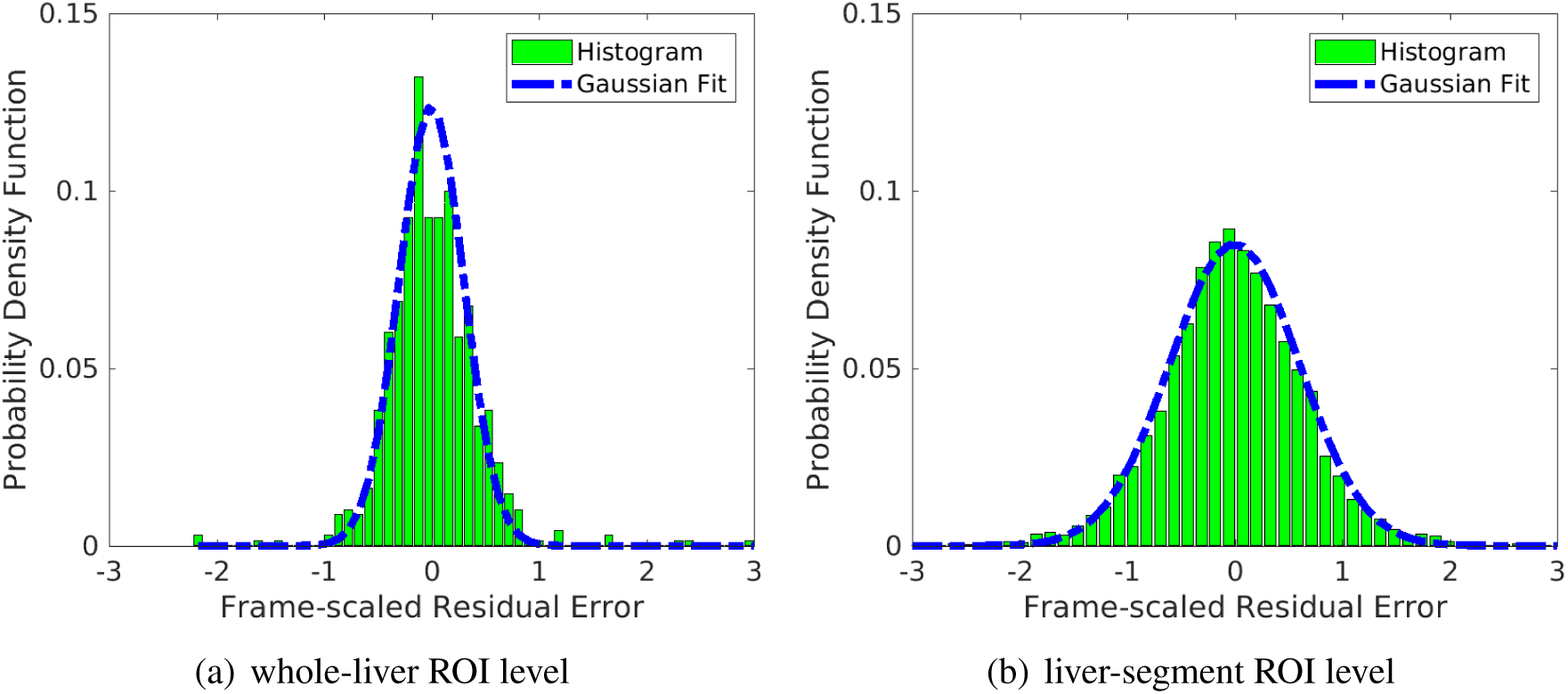
Fit of the histogram of normalized residual difference Δ*c_m_* using a Gaussian distribution with the standard deviation *S_c_*. (a) whole-liver ROI level, *S_c_* = 0.3; (b) liver-segment ROI level, *S_c_* = 0.6.

#### 3.3.3. Comparison of different fitting options

Figure 7 shows the comparison of different initialization and weighting schemes for each patient at the whole-liver ROI level (*S_c_* = 0.3). The *K*_1_ single-initialization strategy resulted in bias in *K*_1_ in several patient datasets. The bias can be reduced when the multi-initialization strategy was used, which however did not provide a universal improvement over all patients. On the other hand, use of nonuniform weighting for TAC fitting led to reduced bias in some patients. The benefit of these two modifications were maximized when they were used together and the bias in *K*_1_ remained small in all patients. Thus, the multi-initialization for *K*_1_ and nonuniform weighting scheme were used in this work for all subsequent analysis.

**Figure 7.**
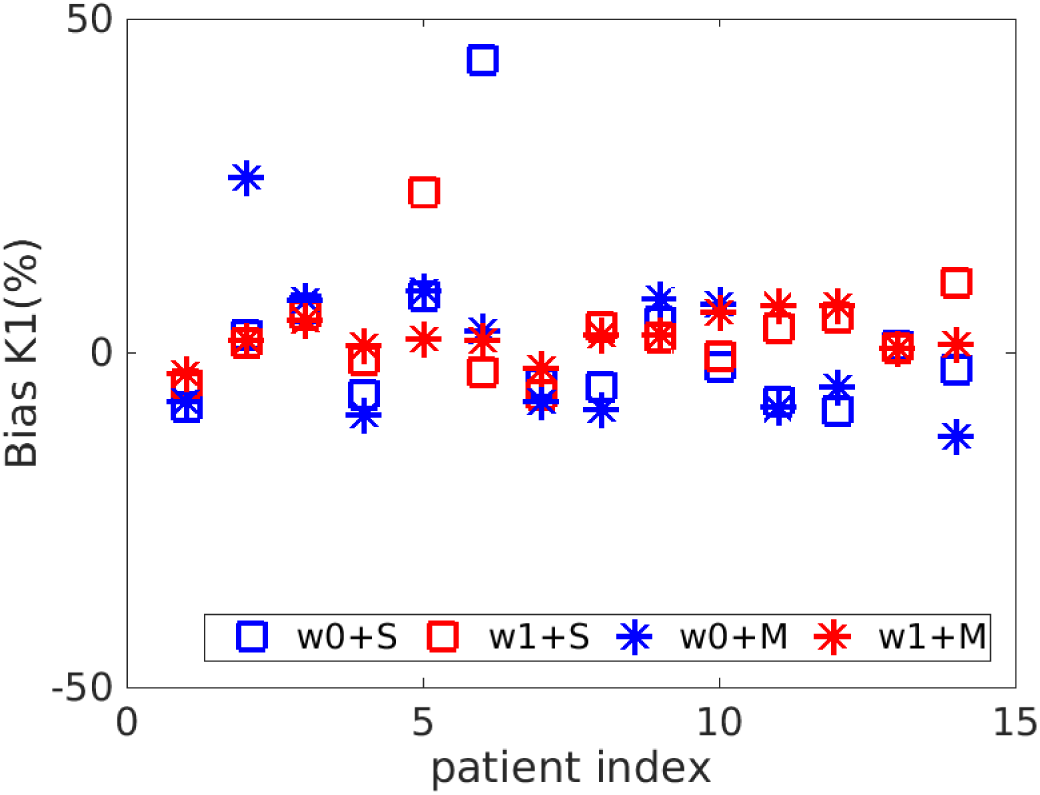
Bias in *K*_1_ estimated using different fitting options. S refers to use single initial value for *K*_1_ and M refers to use multiple initial values. w0 means the uniform weighting scheme and w1 means the nonuniform weighting scheme.

#### 3.3.4. Bias of inappropriate kinetic models

Figure 8 show the bias and SD of *K*_1_ and *K_i_* estimated by three different kinetic models at the noise level of *S_c_* = 0.3. When the traditional SBIF model was used for fitting the TAC which essentially follows the DBIF model, *K*_1_ was underestimated with an average 37% bias. *K*_1_ by the population-based DBIF was also underestimated by 26%. In comparison, the mean absolute bias of *K*_1_ by the optimization-derived DBIF model was only about 3% and the biases of individual patients all remain small. For the estimation of *K_i_*, the SBIF and population-based DBIF resulted in an underestimation of approximately 60%, as compared with an average bias of less than 5% by the optimization- derived DBIF model.

**Figure 8.**
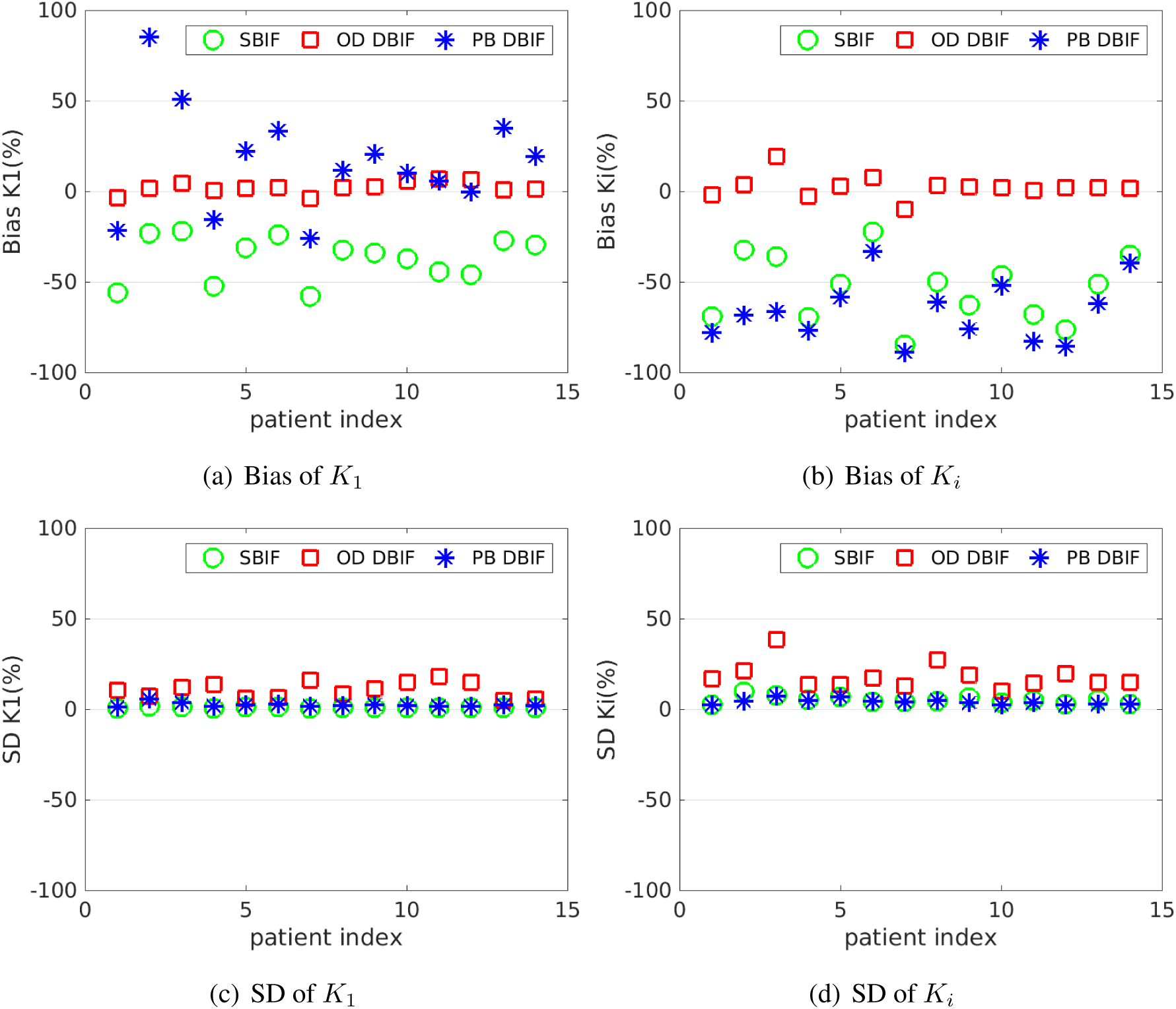
Bias and SD of *K*_1_ and *K_i_* estimates in the optimization-derived (OD) DBIF model for 14 patient data sets with *S_c_* = 0.3, as compared with the inaccurate SBIF model and population-based (PB) DBIF model.

The bias reduction achieved by the optimization-derived DBIF model came with the price of increased SD, as shown in Figure 8(c) and (d). The average SD was 11% for *K*_1_ and 18% for *K_i_* by the optimization-derived DBIF, as compared to less than 6% by the other two models. This can be explained by the increased number of free parameters in the optimization-derived DBIF model.

The results of the averaged absolute bias and SD across different patients are summarized in table 1 for all FDG transport rate parameters. Generally, with the assumption of the TACs following the DBIF model, the inaccurate SBIF model resulted in greater than 35-85% bias and the population-based DBIF model led to greater than 20-90% bias in all kinetic estimates. The accurate optimization-derived DBIF model still had a bias of about 3-8% in all the kinetic estimates, which can be explained by the facts that noise propagation is highly nonlinear and the structural identifiability of the model is local.

**Table 1.** Absolute bias and SD of liver FDG kinetic parameter estimates by different kinetic models. The absolute bias and SD are averaged over 14 patient data sets.

|  | OD DBIF |  | SBIF |  | PB DBIF |  |
| --- | --- | --- | --- | --- | --- | --- |
|  | Bias (%) | SD(%) | Bias(%) | SD(%) | Bias(%) | SD(%) |
| $K_1$ | 3.2 | 10.8 | 36.7 | 1.1 | 25.5 | 2.6 |
| $k_2$ | 3.5 | 11.6 | 40.1 | 1.1 | 21.7 | 2.4 |
| $k_3$ | 5.0 | 19.7 | 56.1 | 5.2 | 69.4 | 4.0 |
| $k_4$ | 8.1 | 44.0 | 85.1 | 9.4 | 90.7 | 8.5 |
| $K_i$ | 4.5 | 18.3 | 53.8 | 5.2 | 66.1 | 4.2 |

Note that as compared to the SBIF and population-based DBIF models, the increase of SD by the optimization-derived DBIF model was generally smaller than the corresponding bias reduction. This indicates the overall gain of the new model is greater than its loss, which led to the improvement in correlating FDG *K*_1_ with histology as we observed in the previous patient study [Wang et al., 2018].

#### 3.3.5. Effect of noise levels on kinetic quantification

Table 2 shows the average absolute bias and SD of kinetic parameters estimated by the optimization-derived DBIF model under three noise levels: noise-free (*S_c_* = 0.0), noise at the whole-liver ROI level (*S_c_* = 0.3), and noise at the liver-segment ROI level (*S_c_* = 0.6). While other kinetic parameters had a small bias, the bias of *υ_b_* and *f_A_* were surprisingly large even at the noise-free case (*S_c_* = 0.0). This can be explained by the fact that the model is locally identifiable with potential multiple solutions. The result is also consistent with the observation on the indifferentiable sensitivity curves of *υ_b_* and *f_A_* in figure 5. Despite the large bias in *υ_b_* and *f_A_*, the bias of *K*_1_ remained small (<8%). The SD of *K*_1_ increased from 7% to 18% when the noise level was changed from the whole-liver ROI level to liver-segment ROI level. The estimation of *K_i_* is more sensitive to noise, with the SD being 18% for the whole-liver ROI level and 37% for the liver-segment ROI level. *k*_2_ had similar accuracy and precision as *K*_1_ and *k*_3_ had similar accuracy and precision as *K_i_*. *k*_4_ had a much higher bias and SD because the scan time (1-hour in our study) is not sufficient enough for robust estimation of *k*_4_ which can justified from its sensitivity curve.

**Table 2.** Bias and SD of the kinetic parameters in the optimization-derived DBIF model under three different noise levels: *S*_c_ = 0 (noise-free), *S_c_* = 0.3 (whole-liver ROI level), and *S_c_* = 0.6 (liver-segment ROI level).

| | $S_c = 0$ | | $S_c = 0.3$ | | $S_c = 0.6$ | |
| --- | --- | --- | --- | --- | --- | --- |
|  | Bias (%) | SD(%) | Bias(%) | SD(%) | Bias(%) | SD(%) |
| $K_1$ | 0.6 | 0 | 3.2 | 10.8 | 7.4 | 18.3 |
| $k_2$ | 0.6 | 0 | 3.5 | 11.6 | 8.2 | 19.6 |
| $k_3$ | 0.2 | 0 | 5.0 | 19.7 | 12.4 | 40.3 |
| $k_4$ | 0.1 | 0 | 8.1 | 44.0 | 22.7 | 81.6 |
| $K_i$ | 0.2 | 0 | 4.5 | 18.3 | 11.1 | 37.0 |
| $v_b$ | 16.2 | 0 | 71.8 | 206.5 | 182.6 | 484.6 |
| $k_a$ | 0.6 | 0 | 3.9 | 16.2 | 6.1 | 27.6 |
| $f_A$ | 16.9 | 0 | 38.3 | 121.9 | 41.6 | 142.5 |

#### 3.3.6. Parameter estimation accuracy over clinical range

Figure 9 shows the plots of linear correlation between the true values and noisy estimates of different kinetic parameters at the whole-liver ROI noise level. The correlation coefficients under different noise levels for all kinetic parameters are summarized in table 3. As the noise level increased, the correlation coefficients reduced. Both *K*_1_ and *K_i_* were well repeatable against noise. While all other kinetic parameters including (*k*_2_, *k*_3_,*k*_4_, *K_a_*) can be repeated well, the two vascular parameters *υ_b_* and *f_A_* are less repeatable.

**Figure 9.**
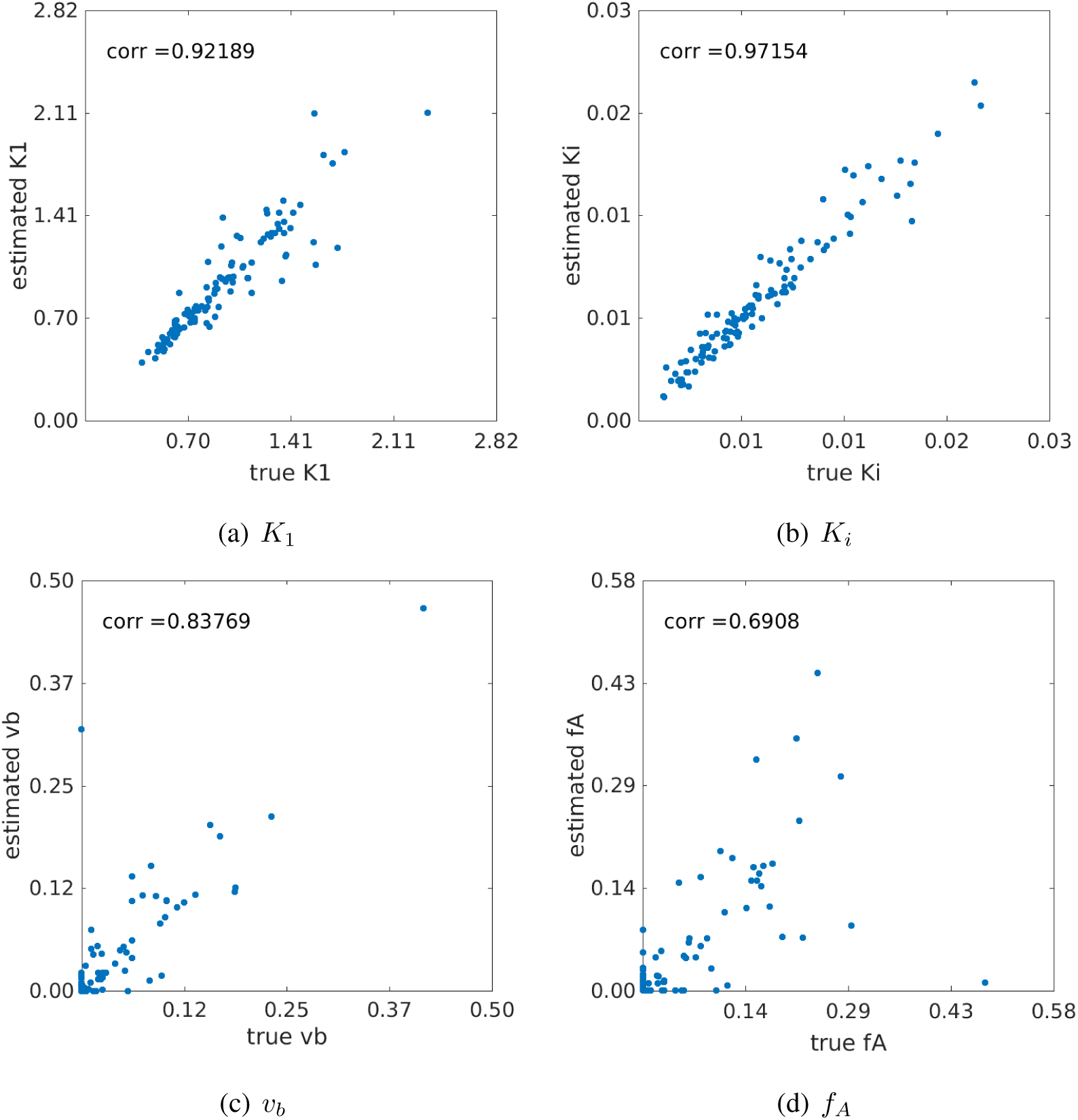
Correlation between the true kinetic parameter values and estimated values from noisy data (*S_c_* = 0.3). (a) *K*_1_, (b) *K_i_*, (c) *υ_b_* and *f_A_*.

**Table 3.** Coefficients of the linear correlation between estimated kinetic parameters and their true values.

| | $v_b$ | $K_1$ | $k_2$ | $k_3$ | $k_4$ | $k_a$ | $f_A$ | $K_i$ |
| --- | --- | --- | --- | --- | --- | --- | --- | --- |
| $S_c = 0.3$ | 0.84 | 0.92 | 0.90 | 0.97 | 0.98 | 0.91 | 0.69 | 0.97 |
| $S_c = 0.6$ | 0.78 | 0.88 | 0.83 | 0.90 | 0.91 | 0.79 | 0.66 | 0.90 |

#### 3.3.7. Noise variation of the *K*_1_ correlation with liver inflammation

Figure 10 shows the results of correlating the histological inflammation scores with the FDG *K*_1_ estimates derived from 1000 noisy realizations (*S_c_* = 0.3). The linear correlation *r* between the original *K*_1_ values and liver inflammation scores in the cohort of 14 patients was *r* =-0.7618 (p=0.0012). The 95% confidence interval of the noisy *r* estimates was estimated to be [-0.8434, −0.6494] with the mean −0.7452 and standard derivation 0.0493. The percent bias in *r* was −2.2% and the SD was 6.5%, both approximately close to that in *K*_1_. These results indicate the stability of the estimation of the correlation between FDG *K*_1_ and liver inflammation.

**Figure 10.**
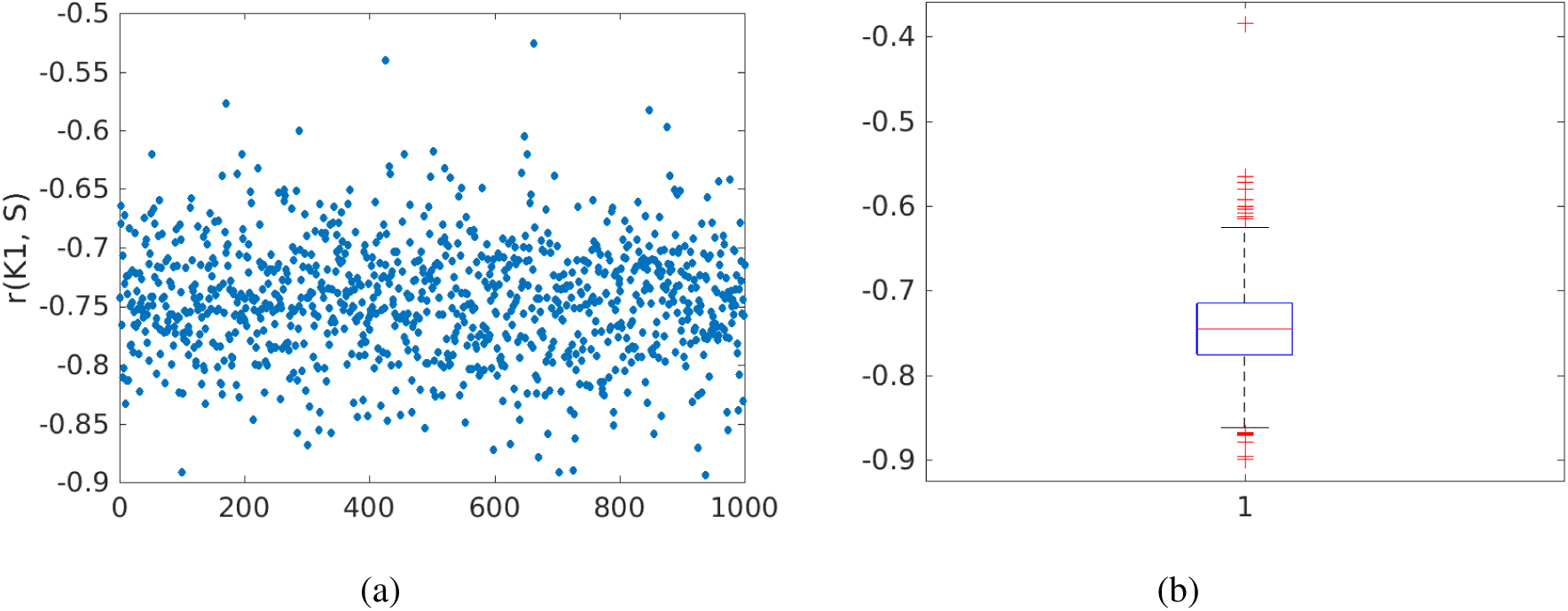
Noise variation of the correlation *r* between FDG *K*_1_ and histological liver inflammation score. (a) *r* values of 1000 noisy realizations, (b) box plot of the *r* values.

## 4. Discussion

FDG *K*_1_ by the optimization-derived DBIF model is a promising PET biomarker for evaluating human liver inflammation in fatty liver disease [Wang et al., 2017, Sarkar et al., 2017, Wang et al., 2018]. The focus of this work is to characterize the identifiability of the optimization-derived DBIF model structure and evaluate the accuracy and precision of *K*_1_ and other kinetic parameters in dynamic liver PET.

We first conducted a theoretical analysis of the structural identifiability of standard 3-compartmental model and the new DBIF model using the Laplacian transform. While standard 3-comaprtmental model is globally identifiable, the new model is locally identifiable due to potential multiple solutions. This suggests that it is worth being careful with defining the initial values, lower and upper bounds of kinetic parameter estimation to properly constrain the optimization problem of TAC fitting with the new model.

We then conducted Monte Carlo simulations to examine the practical identifiability of the model parameters based on 14 patient datasets which include both dynamic FDG-PET data and histopathology data of human liver inflammation. While the estimation of some kinetic parameters (e.g. *f_A_* and *υ_b_*) is associated with large bias and standard deviation, FDG Ki, the parameter of major interest, has low bias (≈3%) and standard deviation (≈11%) at the whole-liver ROI level. As demonstrated in the simulation study, fitting liver TACs using the traditional SBIF model or the population-based DBIF model may result in significant bias (>20%) in liver *K*_1_ quantification. These results explain why the *K*_1_ by the new model achieved a statistically significant association with liver inflammation in the patient study, while the other two models did not demonstrate success [Wang et al., 2018].

We also examined the reliability of the new model for liver *K*_1_ quantification over a wide range of values from 0.5 to 2.5 (Fig. 9). The true *K*_1_ values and their estimates are highly correlated (*r* > 0.9). The stability of *K*_1_ estimation against noise is also preserved in its correlation with liver inflammation (Fig. 10).

A disadvantage of the optimization-derived DBIF model is the increased standard deviation in kinetic parameter estimation, which is basically caused by the increased number of free parameters. To control the standard deviation of *K*_1_ and other parameters of interest, one potential strategy is to add additional constraints in the optimization problem. For example, Table 4 compares the bias and standard deviation of kinetic parameters for either estimating or fixing the input function parameter *k_a_* in the optimization of TAC fitting. If *k_a_* is fixed at its true values, the bias and standard deviation of *K*_1_ (and other kinetic parameters) can be largely reduced. This is not surprising because a fixed *k_a_* corresponds to a known portal vein input function. However, the result reported here indicates the potential improvement space if a modified method can be developed to incorporate the prior information of the portal vein input function.

**Table 4.** Bias and SD of a typical kinetic parameter set (population means) estimated by the optimization-derived DBIF model with *k_a_* freely estimated or fixed at its true value.

| | free $k_a$ | | fixed $k_a$ | |
| --- | --- | --- | --- | --- |
|  | Bias (%) | SD(%) | Bias(%) | SD(%) |
| $K_1$ | 3.2 | 10.8 | 0.3 | 3.0 |
| $k_2$ | 3.5 | 11.6 | 0.4 | 3.4 |
| $k_3$ | 5.0 | 19.7 | 1.9 | 17.7 |
| $k_4$ | 8.1 | 44.0 | 4.6 | 41.6 |
| $K_i$ | 4.5 | 18.3 | 1.7 | 16.6 |
| $v_b$ | 71.8 | 206.5 | 47.9 | 116.9 |
| $k_a$ | 3.9 | 16.2 | / | / |
| $f_A$ | 38.3 | 121.9 | 40.0 | 106.0 |

The study also indicates that kinetic quantification at the liver-segment ROI noise level (*S_c_* = 0.6) is less reliable than at the whole-liver ROI noise level (*S_c_* = 0.3). Both bias and SD become nearly doubled, as shown in Table 2. It is worth noting that all the studies were conducted using standard clinical PET. Recently, high-sensitivity total-body PET scanner EXPLORER has been developed [Cherry et al., 2018]. The new generation scanner can increase sensitivity of PET by a factor of 4-5 for imaging a single organ [Poon et al., 2012]. The scanner sensitivity improvement will be able to reduce the liver-segment ROI noise level from current *S_c_* = 0.6 to *S_c_* = 0.3. Thus, quantification of liver segmental heterogeneity may become reliable on EXPLORER. Equivalently, the whole-liver ROI noise level may also be reduced from current *S_c_* = 0.3 to *S_c_* = 0.15 if EXPLORER is used. The resulting bias and SD of *K*_1_ were 1.4% and 6.5%, respectively, according to our simulation study.

## 5. Conclusion

This paper has conducted both theoretical analysis of structural identifiability and Monte Carlo study of practical identifiability for the optimization-derived DBIF model in dynamic PET of liver inflammation. The theoretical analysis suggests that the parameters of the new model are identifiable but subject to local solutions. The simulation results have shown that the estimation of vascular kinetic parameters (*υ_b_* and *f_A_*) suffer from high variation. However, FDG *K*_1_ can be reliably estimated in the new optimization-derived DBIF model. The bias of *K*_1_ by the new model is approximately 3% and the standard deviation is about 11% at the whole-liver ROI noise level. The estimated values and original values of *K*_1_ are also highly correlated with each (*r* = 0.92). The correlation between liver *K*_1_ by the new model and histological inflammation score is robust to noise interference. These results suggest that liver FDG *K*_1_ quantification is reliable for clinical use to assess liver inflammation at the whole-liver ROI level. Future work will include further development of the DBIF modeling approach and use of EXPLORER for reduced bias and variance in *K*_1_.

